# Fast and easy single-molecule pulldown assay based on agarose microbeads

**DOI:** 10.1101/2020.09.20.305177

**Authors:** Qirui Zhao, Yusheng Shen, Xiaofen Li, Fang Tian, Xiaojie Yu, Levent Yobas, Hyokeun Park, Pingbo Huang

## Abstract

The recently developed single-molecule pulldown (SiMPull) assay by Jain and colleagues is a highly innovative technique but its wide application is hindered by the high technical barrier and time consumption. We report an innovative, agarose microbead-based approach for SiMPull. We used commercially available, pre-surface-functionalized agarose microbeads to capture the protein of interest together with its binding partners specifically from cell extracts and observed these interactions under a microscope at the single-molecule level. Relative to the original method, microbead-based SiMPull is considerably faster, easier to use, and more reproducible and yet provides similar sensitivity and signal-to-noise ratio; specifically, with the new method, sample-preparation time is substantially decreased (from ∼10 to ∼3 h). These crucial features should facilitate wide application of powerful and versatile SiMPull in common biological and clinical laboratories. Notably, by exploiting the simplicity and ultrahigh sensitivity of microbead-based SiMPull, we used this method in the study of rare auditory hair cells for the first time.

## INTRODUCTION

A protein never works alone: it works together with other proteins that serve as its regulators, auxiliary subunits, or effectors. The functional link between two proteins typically results from their intimate physical interaction, or protein-protein interaction (PPI), although the link is occasionally mediated by soluble factors such as cAMP or Ca^2+^. Therefore, PPIs are indispensable for nearly all aspects of diverse cellular processes, and assessment of the PPIs of a protein of interest is essential for elucidating the function and regulation of the protein.

For studying PPIs in general, a powerful and commonly used technique is conventional coimmunoprecipitation (co-IP/pulldown) followed by western blotting. However, the technique does not provide precise information regarding the kinetics and stoichiometry of PPIs. Another drawback is that conventional co-IP cannot be used for examining PPIs in rare cells such as sensory hair cells, circulating/residual tumor cells, embryonic stem cells, and subsets of immune cells: A single western blotting assay and a single co-IP assay typically require ∼10^4^–10^5^ cells (Schulte et al., 2008) and ∼10^5^–10^6^ cells, respectively, and rare cells cannot be readily acquired in such large quantities. This technical constraint in conventional co-IP substantially retards research on rare cells such as sensory hair cells.

Recently, Jain et al. developed a single-molecule pulldown (SiMPull) assay combining the principles of conventional pulldown and single-molecule fluorescence microscopy (Jain et al., 2011). The highly innovative SiMPull method is superior to the conventional co-IP assay in several respects: SiMPull enables not only stoichiometric and kinetic evaluation of a protein complex, but also examination of relatively weak and transient PPIs featuring Kd values in the micromolar range(Lee et al., 2013), which are readily disrupted during the multiple washings employed in conventional co-IP. Moreover, because of its ultrahigh sensitivity, SiMPull can be used to analyze PPIs in as few as 10 cells (Jain et al., 2011) and can thus potentially be used for studying rare cells. However, SiMPull has not been widely used since its development, which is probably because of the high technical barrier and time consumption of the method.

Here, we report an innovative agarose microbead-based approach for SiMPull. Our new method is considerably simpler and faster than the original SiMPull of Jain et al., and these crucial features should greatly facilitate wide application of powerful SiMPull in common biological and clinical laboratories. Moreover, we tested the microbead-based SiMPull assay in the study of rare hair cells for the first time.

## METHODS AND MATERIALS

### Materials

The following chemicals and other materials were from commercial sources: methanol [Catalog (Cat.) # BDH1135], acetone (Cat.# BDH1101), and 2-propanol (Cat.# BDH1133), BDH Chemicals; PEI 25000 (Cat.# 23966-1), Polysciences; Tris-HCl (Cat.# BP153-1), Fisher Scientific; PBS (Cat.# 10010-023), Gibco; sodium deoxycholate (Cat.# D6750) and NP-40 (Cat.# N3500), United States Biological; biotin-Atto 488 (Cat.# 30574) and bovine serum albumin (BSA; Cat.# A7030), Sigma-Aldrich; NeutrAvidin (Cat.# 31000), Pierce; NeutrAvidin agarose microbeads (Cat.# 29201), Thermo Scientific; quartz slides (Cat.# 7101), Sail Brand; and coverslips (24×24 mm, Cat.# 48393230), VWR International.

### Antibodies

These antibodies were purchased: sheep polyclonal anti-PCDH15 (Cat.# AF6279; R&D Systems); mouse monoclonal anti-HA (MMS-101p; Covance) and anti-FLAG (clone M2; Sigma-Aldrich); Alexa Fluor 488-conjugated (Cat.# ab181448; Abcam) and Alexa Fluor 647-conjugated (Cat.# ab150115; Abcam) goat anti-mouse IgG; biotinylated (Cat.# 65-6140; Thermo Fisher) and Alexa Fluor 647-conjugated (Cat.# ab150079; Abcam) goat anti-rabbit IgG; Alexa Fluor 488-conjugated donkey anti-sheep IgG (Cat.#ab150177); and biotinylated donkey anti-rabbit IgG (Cat.# A16027; Thermo Fisher).

The following homemade antibodies were used in this study: rabbit anti-TMC1 serum (against N-terminal 39 aa residues of human TMC1)(Li et al., 2019), anti-GFP serum (against purified GFP), and anti-LHFPL5 serum (against C-terminal 20 aa residues of human LHFPL5; Yu et al., Under Revision). The antibodies were generated by immunizing rabbits housed in the animal care facility at the Hong Kong University of Science and Technology.

### Solutions

T50-BSA buffer contained 50 mM NaCl, 10 mM Tris-HCl, 0.1 mg/mL BSA, pH 7.5 adjusted with HCl, and the wash buffer was T50-BSA buffer without BSA; both solutions can be stored at 4°C for up to 1 month. The lysis buffer contained 150 mM NaCl, 10 mM Tris, 1% (v/v) NP-40, 1 mM EDTA, protease inhibitors (cOmplete mini, ROCHE), pH 7.5 adjusted with HCl; the buffer was freshly prepared for each use.

### Cell culture, transfection, and expression vectors

The procedures used for cell culture and transfection were almost identical to those published previously (Hu et al., 2017). HEK293T cells (RRID: CVCL_1926) were obtained from American Type Culture Collection (ATCC, Manassas, VA, USA); the cells were presumably authenticated by ATCC and were not further authenticated in this study. The cell line was routinely confirmed to test negative for mycoplasma contamination. HEK293T cells were maintained in Dulbecco’s modified Eagle medium supplemented with 10% fetal bovine serum and 100 U/mL penicillin/streptomycin (Life Technologies) in an atmosphere of 95% air–5% CO2 at 37°C. Cells were transfected using polyethylenimine (3 μL/μg plasmid).

The expression vectors used in the study were pEGFP-n1 for GFP expression (Cat.# 6085-1), pcDNA3-mouse PKA-Calpha-mEGFP (Cat.# 45521), and pcDNA3-mouse PKA-RIIalpha-mEGFP (Cat.# 45527), all from Addgene. pcDNA3-PKA-Calpha-mCherry was constructed by replacing the GFP sequence in pcDNA3-PKA-Calpha-mEGFP with mCherry sequence.

### Preparation of primary antibody-coated agarose microbeads

All procedures were performed at room temperature. Briefly, 0.5 μL of NeutrAvidin-coated agarose microbeads (bed volume ∼50%) were incubated with 100 μL of 10 nM biotinylated 2^nd^ antibody in PBS for 10 min, after which the microbeads were spun down by centrifugation at 150 × *g* for 90 s in Eppendorf tubes and washed twice with PBS; this centrifugation step removed some of the smaller microbeads as well. Subsequently, the microbeads were incubated with 100 μL of 1^st^ antibody against the bait protein in T50-BSA buffer (1:100 anti-TMC1 or anti-LHFPL5 serum, or 1:50 anti-GFP) for 20 min, washed thrice with PBS, and stored in 100 μL of PBS until use (We recommend the use of the beads within 1 h at room temperature). Between solution changes, microbeads were spun down at 150 × *g* for 90 s in these and following procedures.

### Protein pulldown using agarose microbeads from cultured cells

In an Eppendorf tube, ∼10^5^ cells were lysed with 100 μL of lysis buffer, and then 10 μL of primary antibody-coated microbeads were mixed with the cell lysate and incubated for 30 min. After washing with 200 μL of wash buffer twice, the microbeads were resuspended in wash buffer at a final volume of 10 μL. Subsequently, the microbeads were transferred onto a glass slide and covered with a coverslip (18×18 mm). The coverslip was precleaned through sonication in acetone, isopropanol, and water (5 min each). Because the prey proteins were fluorescently tagged in these experiments, the proteins captured on the microbeads could be imaged immediately under a total internal reflection fluorescence (TIRF) microscope or a confocal microscope.

### Protein pulldown using agarose microbeads from primary cells (organ of Corti cells)

Mice were euthanized by decapitation and then entire cochleae were dissected in Hank’s Balanced Salt Solution containing 0.1 mM CaCl2. The organ of Corti was further dissected from the cochlea and the tectorial membrane was removed, and the tissue samples (2–3 organs of Corti) were transferred into 60 μL of lysis buffer, ground thoroughly in a mini tissue grinder (Cat.# 357848, Wheaton), and sonicated for 2 min on ice. Subsequently, tissue lysates were centrifuged for 15 min at 13,000 × *g* to remove cell debris and then incubated with 10 μL of 1^st^ antibody-coated microbeads in PBS at room temperature for 30 min.

Because the prey proteins in these experiments were not fluorescently tagged, 100 μL of the prey antibody in T50-BSA buffer (1:200 anti-FLAG or 1:50 anti-PCDH15) was added to the Eppendorf tubes containing the samples and incubated for 30 min. After washing with 200 μL of wash buffer thrice, 50 μL of 20 nM fluorophore-labeled 2^nd^ antibody in T50-BSA buffer was added to the tubes and incubated for 15 min, and after washing thrice more with the wash buffer to remove unbound antibodies, the microbeads were resuspended in a final volume of 10 μL of wash buffer and examined under a microscope (following procedures similar to those used for cultured cells).

### Single-molecule imaging and photobleaching under a TIRF microscope

Fluorescence images were taken using an Olympus IX-73 inverted fluorescence microscope (Olympus) equipped with an oil-immersion objective (NA = 1.49, 100×, UAPON, Olympus) as previously described (Alsina et al., 2017) with minor modifications. Images were acquired using an sCMOS camera (Zyla-4.2P-Cl10, Andor Technology Ltd.). A 488-nm laser (Coherent Inc., USA) was used to excite GFP with an exposure time of 300 ms. Only GFP near the surface of coverslips were excited using an objective total internal reflection illumination. ZT488/561rpc (Chroma) was used as a dichroic mirror, and the emission signals were collected through an ET525/50m (Chroma) emission filter to collect fluorescent signals from GFP. The lasers were focused on the back focusing plane of the objective. A 15X beam expander (Edmond optics, Singapore) and a focus lens were used to illuminate the sample uniformly. Total internal reflection illumination was produced by moving the illumination beam away from the center of the lens. All experiments were performed at room temperature.

In photobleaching experiments, the fluorescence time-traces of GFP molecules on agarose microbeads were acquired with a 100-s time span and 2-Hz frame rate (an exposure time of 0.3 s). We first selected fluorescent spots whose intensity profiles displayed a Gaussian distribution with a sigma of <130 nm, and the selected spots (∼80% of total spots) were subject to photobleaching analysis. The photobleaching trace and steps were manually determined using ImageJ software, following published procedures (Ulbrich & Isacoff, 2007). Fluorescent spots were further discarded in the bleaching analysis if their trace did not display discrete decreasing steps, if their intensity did not drop to the background level, or if they blinked in 5 consecutive frames. Subsequently, fluorescent spots were classified as exhibiting 1, 2, or >2 bleaching steps.

### Single-molecule imaging under a confocal microscope

The confocal microscope used in the study was a Leica TCS SP8 Confocal Microscope. Excitation lasers at 488 nm, 514 nm, 552 nm, 638 nm were generated by diode generators, and a 63× objective lens (Leica HC PL APO) was used to examine samples; the fluorescent signal was collected using the same objective and imaged by using an HyD SP GaAsP detector. GFP and mCherry were excited (at a scanning speed of 100 Hz) at 488 nm with 1% maximum laser power and 552 nm with 3% maximum laser power, respectively.

### Statistics

All data are expressed as means ± SEM; n denotes the number of independent biological replicates. Unless indicated otherwise, Student’s two-tailed *t* test was used for statistical analysis, and P < 0.05 was considered statistically significant.

## RESULTS

### Microbead-based strategy substantially simplifies SiMPull sample preparation

The SiMPull technique developed by Jain et al. enables PPI detection at the single-molecule level and thereby offers several advantages over other methods (see “Introduction”)(Jain et al., 2011). However, the technique requires meticulous multistep cleaning, surface-passivation, and functionalization of quartz slides, and these procedures are extremely time-consuming and prone to large variations (Supplementary Table 1). Our hands-on experience suggests that this represents one of the major hurdles in the wide application of the powerful SiMPull technique in common biological research and clinical laboratories.

To overcome the aforementioned drawbacks, we used commercially available, pre-surface-functionalized agarose microbeads (Fig. 1 and Supplementary Table 1). Our microbead-based method not only considerably saves time by omitting quartz-slide functionalization, but also offers high reproducibility because the pre-surface-functionalized microbeads are subject to industry-level quality control. Furthermore, the SiMPull method of Jain et al. requires a micro flow-chamber on slides (Jain et al., 2011), and both the preparation of the flow-chamber and the low flow rate of the chamber (which substantially slows the multiple solution changes) make the method highly time-consuming; by contrast, in our microbead-based SiMPull, the flow-chamber is replaced with an Eppendorf tube for all sample-preparation procedures, including bait-antibody coating, lysate incubation, detection with prey antibody, and multiple washes (Fig. 1 and Supplementary Table 1). Collectively, these features of microbead-based SiMPull make this method considerably faster, easier to use, and more reproducible than the original method (Jain et al., 2011) while maintaining similar sensitivity and signal-to-noise (S/N) ratio. Notably, the microbead-based approach reduced sample-preparation time from ∼10 h (Jain et al., 2012) to ∼3 h (Supplementary Table 1).

**Figure 1.**
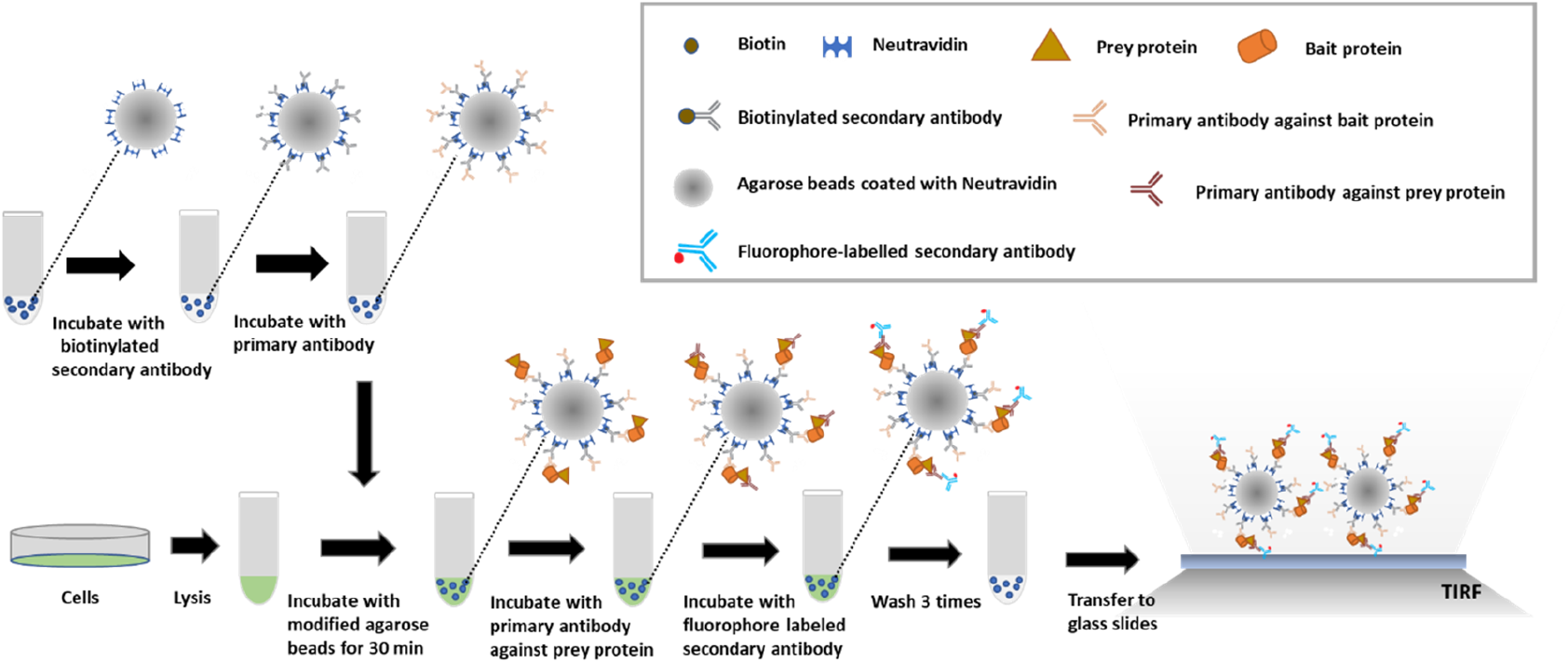
Schematic of agarose microbead-based SiMPull. NeutrAvidin-coated agarose beads are modified by immobilizing a biotinylated 2^nd^ antibody and the corresponding 1^st^ antibody on their surface for capturing the target (bait) protein. Cells are lysed and the agarose microbeads are added to the lysates and incubated for 30 min in an Eppendorf tube. After capture by the microbeads, the prey protein is detected using a specific 1^st^ antibody and a fluorescently labeled 2^nd^ antibody. Microbeads are washed thrice to remove nonspecifically bound proteins, transferred to glass slides, covered with coverslips, and imaged using a TIRF or confocal microscope. Some of the washing steps are omitted in the schematic for the sake of simplicity (see additional details in “Methods and Materials”), and the microbeads are spun down between solution changes. In certain cases, the bait or prey protein is fluorescently labeled and can be visualized directly without immunostaining.

We first evaluated the detection sensitivity of our microbead-based SiMPull method (Supplementary Fig. 1). Biotinylated Atto 488 was used for this test because the extremely high binding affinity of NeutrAvidin for biotin (Kd = 10^−15^ M) ensures that almost all the biotin molecules added at a sub-saturation concentration bind to NeutrAvidin; this feature allows accurate assessment of the upper limit of the detection sensitivity. We found that microbead-based SiMPull was highly sensitive: biotinylated Atto 488 was detected at a concentration as low as 10 pM with an S/N ratio of >10; in these experiments, the negative controls were biotinylated Atto 488 at 100 fold higher concentration (1 nM) with microbeads pre-exposed with biotin, and IgG-conjugated Alexa 488 used at 10,000-fold higher concentration (100 nM) (Supplementary Fig. 1). Moreover, we checked for the nonspecific binding of other proteins such as BSA and ovalbumin to the microbeads, and we again found minimal binding, even after 4 h incubation, because agarose is highly hydrophilic (Supplementary Fig. 2).

**Figure 2.**
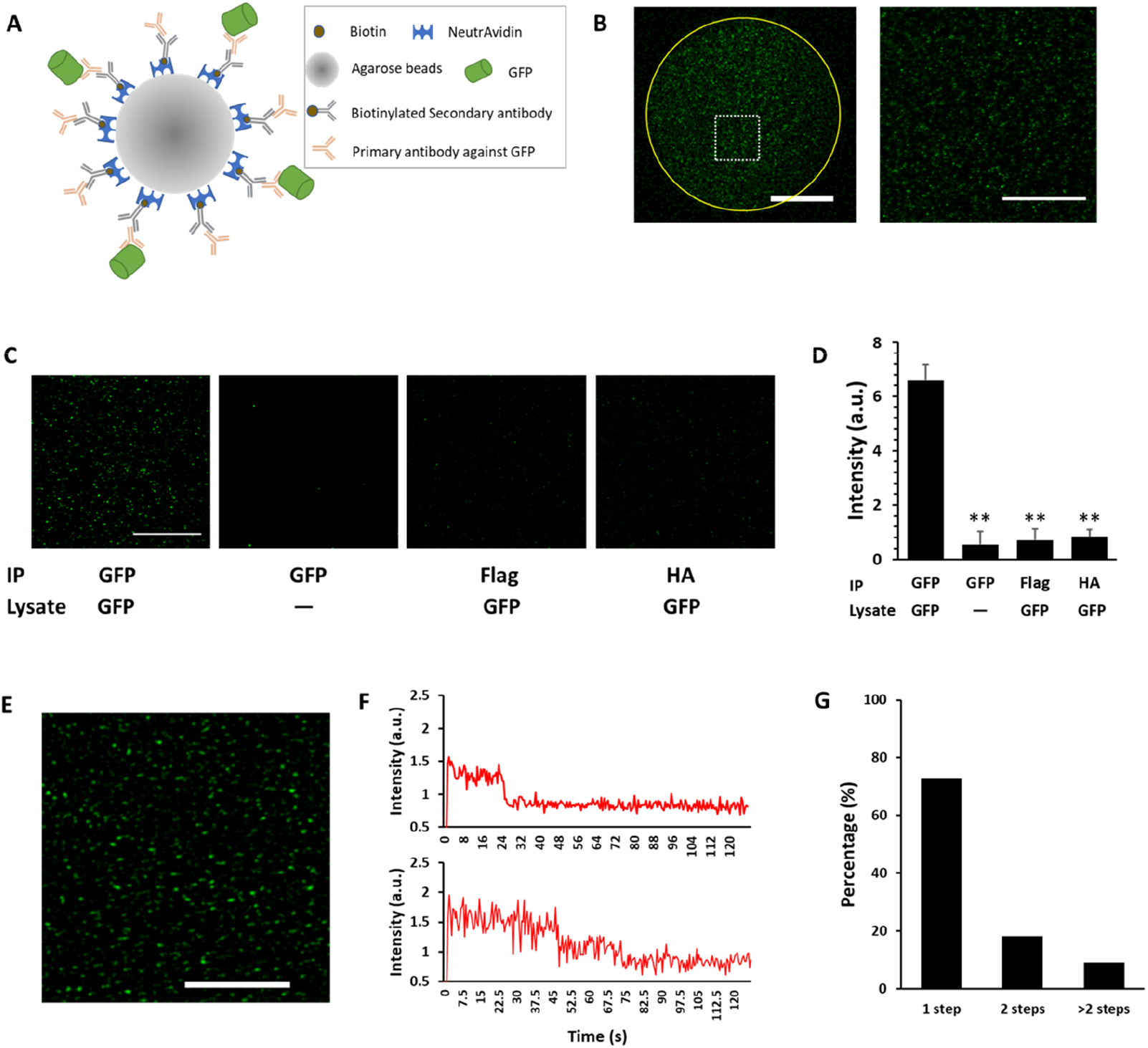
GFP pulldown using microbead-based SiMPull. **A-B)** Schematic of experimental setup (A) and selection of imaging area for analysis (B). GFP was pulled down using NeutrAvidin-coated agarose microbeads that were surface-modified with anti-GFP (A). An imaging area of 25×25 μm (B, right) was selected from a microbead (B, left) for analysis in Panel C. Scale bars, 30 μm (left) and 10 μm (right). **C)** GFP in the pre-made lysate of GFP-expressing or non-transfected HEK293T cells was pulled down using microbeads surface-coated with anti-GFP (see Panel A), anti-FLAG, or anti-HA antibody and then examined under a confocal microscope. The high signal-to-noise (S/N) ratio suggests that GFP was specifically pulled down by anti-GFP antibody. Scale bar, 10 μm. **D)** Statistical results from Panel C and 2 similar experiments. Different from 1^st^ group on the left: \*\**P* ≤ 0.0096; n = 3 independent biological replicates, each representing the average fluorescence intensity of 7 imaging areas in the same experiment. **E-G)** TIRF image (E) of a imaging area from a microbead after GFP pulldown. The spots in Panel E typically displayed one-step (upper) and two-step (lower) bleaching in a photobleaching experiment (F). Panel G: photobleaching-step distribution of 100 selected fluorescent spots from Panel E. Scale bar, 10 μm.

### Microbead-based SiMPull for GFP pulldown

For assessing the performance of microbead-based SiMPull in IPs, we first examined the capture of GFP; this is because GFP can be directly visualized without immunostaining, which simplifies the validation, and, more importantly, because GFP can be used in photobleaching experiments to evaluate the single-molecule state of a fluorescent spot captured on the microbeads.

Ectopic GFP expressed in HEK293T cells was pulled down using microbeads (Fig. 2A-D), whereas few GFP molecules were captured in the 3 negative controls included in the experiments (Fig. 2C-D); the S/N ratio in these experiment was ∼10 (Fig. 2D), which was comparable to that of the original SiMPull assay (Jain et al., 2011).

Furthermore, we performed photobleaching assays on the fluorescent spots after discarding the spots that displayed non-Gaussian distribution and features unexpected for native GFP molecules (see “Methods and Materials”). By using published procedures (Ulbrich & Isacoff, 2007), we found that 73% of the selected spots contained single fluorophores (i.e., GFP monomers), 18% contained 2 fluorophores, and 9% contained >2 fluorophores (Fig. 2E-G). The fluorescent spots that exhibited non-Gaussian distribution probably also represented ≥2 fluorophores located in close proximity, and both this class of spots and the fluorescent spots that displayed Gaussian distribution but contained ≥2 fluorophores were likely generated due to the high density of GFP molecules on the microbead surface; conceivably, the proportion of both of these classes of fluorescent spots can be reduced by diluting antibodies or GFP molecules. Nevertheless, our results clearly demonstrated that the microbead-based approach enables a protein of interest to be trapped at the single-molecule level at a very high S/N ratio, and this represents one of the powerful features of the SiMPull method (Jain et al., 2011).

### Signal homogeneity and minimal cell number for protein pulldown

The microbeads used in this study displayed inherent size variation and were predominantly 40–70 μm in diameter (Supplementary Fig. 3). For the data analysis in the assay in Fig. 2C-D and in other similar experiments, we used the average signal intensity of a small number (6–7) of randomly selected microbeads as a faithful representation of all the microbeads in the same experiment; this approach was based on the assumption that the microbead-surface density of the captured bait protein was homogenous and independent of bead size. We used GFP pulldown experiments to verify this assumption. As per our expectation, the counts of the fluorescent dots and the total fluorescence intensity per surface area were extremely close (Fig. 3A-D). A bright aureole was detected at the edge of the microbeads (Fig. 3A) and was presumably generated from the curved edge of the beads (Supplementary Fig. 4); however, the aureole was invisible in images captured at high magnification and low optical depth (e.g., Figs. 2E and 4B).

**Figure 3.**
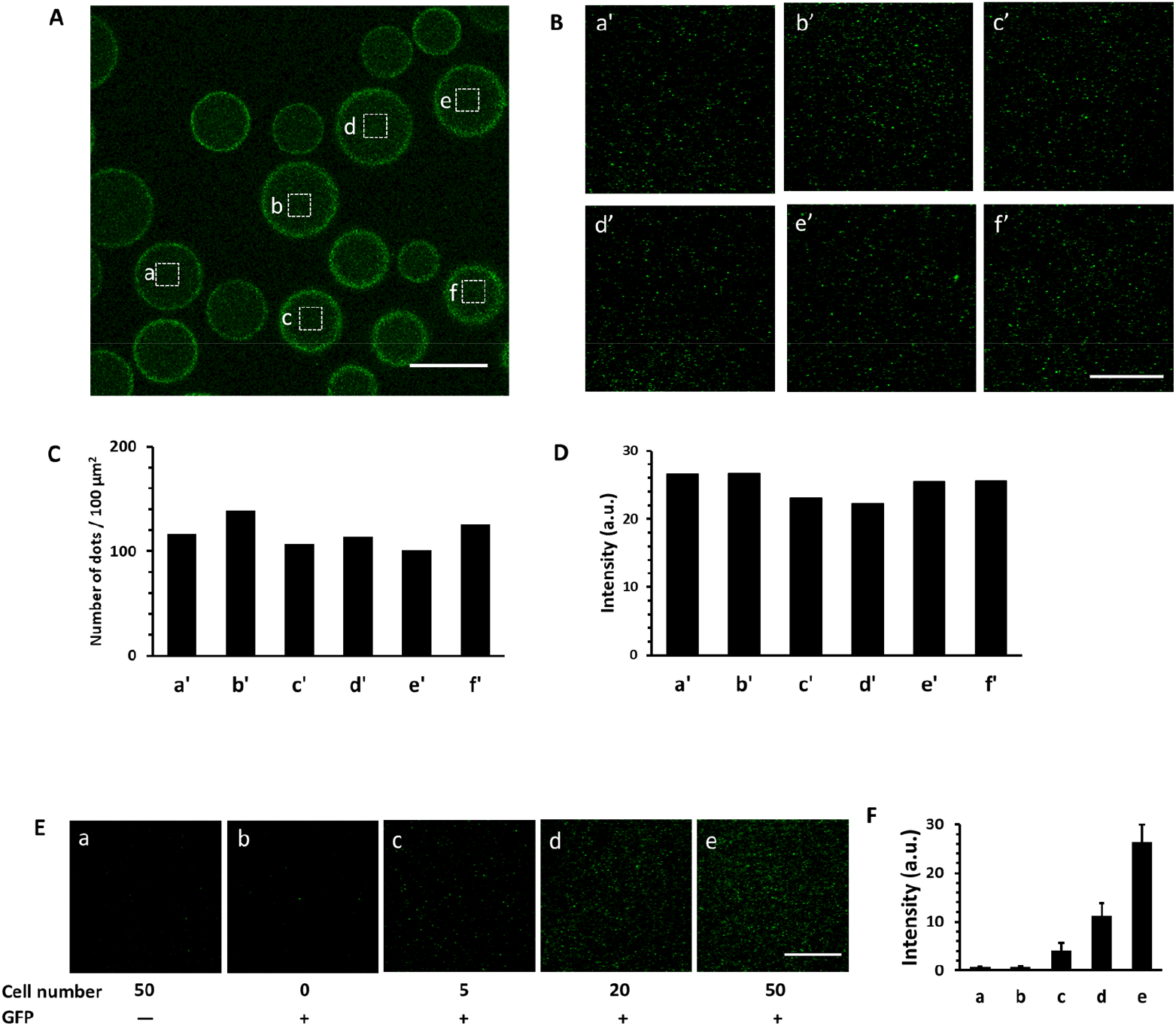
Signal homogeneity and minimal cell number of microbead-based SiMPull. **A-B)** GFP in the pre-made lysate of GFP-expressing HEK293T cells was pulled down using anti-GFP surface-coated NeutrAvidin microbeads. Fluorescent spots in a 25×25 μm imaging area were sampled from 6 randomly selected microbeads of various sizes (A). Panel B: magnification of boxed imaging areas in Panel A. **C-D)** Counts of fluorescent dots (C) and total signal intensity (D) of each imaging area in Panel B. **E-F)** GFP-expressing HEK293T cells were lysed and diluted at distinct ratios to represent various cell numbers and then subject to microbead-based SiMPull for GFP. Fifty non-transfected HEK293T cells are used as a negative control (a). Panel F: statistical results from Panel E and 2 similar independent biological replicates, each representing the average fluorescence intensity of 7 imaging areas in the same experiment. The S/N ratios are calculated from the signal intensities by using the signal intensity of 50 cells without GFP as noise background: 6.3 (5 cells), 17.2 (20 cells), and 40.8 (50 cells). Scale bars, 100 μm in Panel A and 10 μm in Panels B and E.

**Figure 4.**
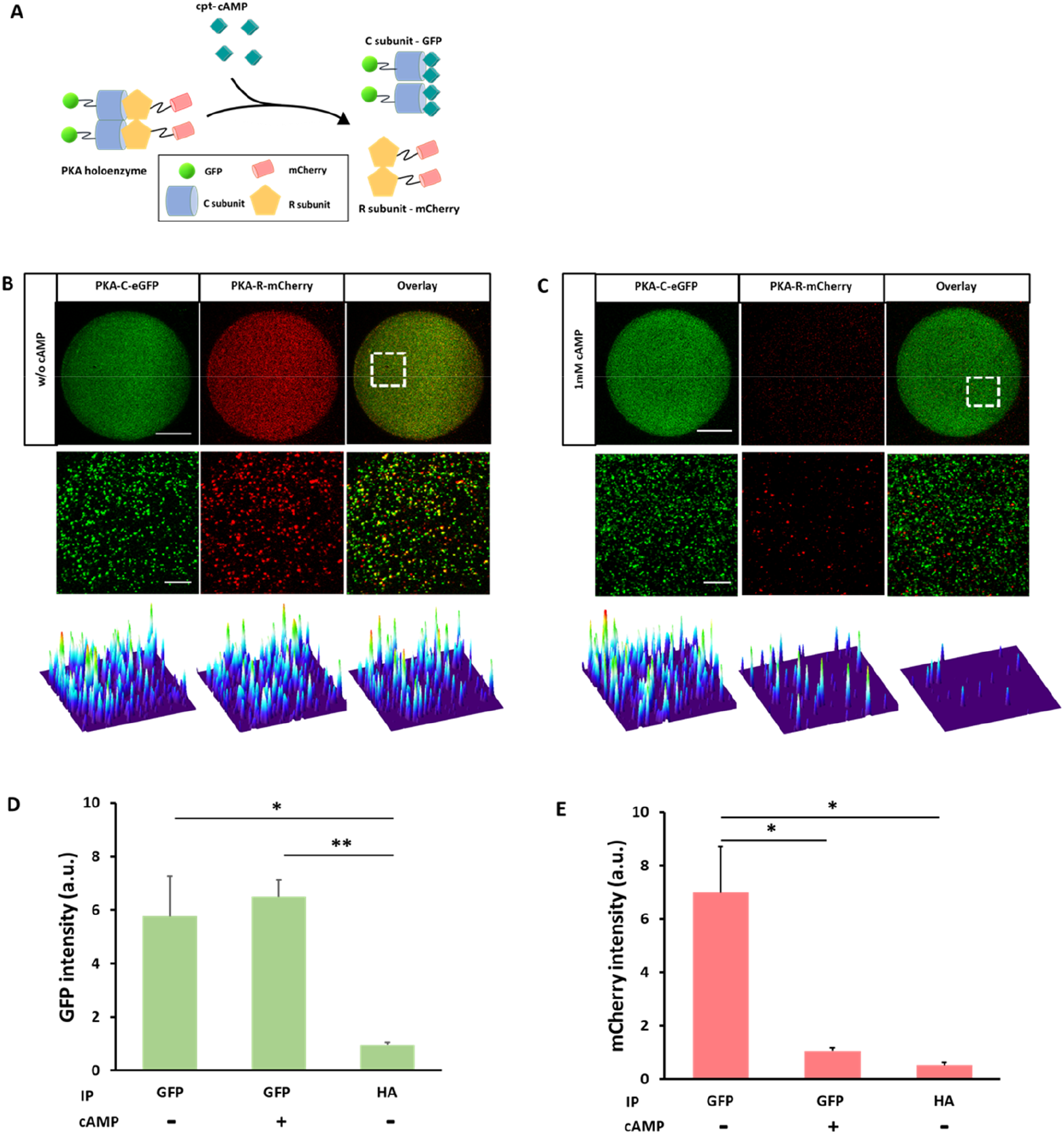
Pulldown of PKA complex by using microbead-based SiMPull. **A)** Schematic of PKA complex and its activation by cAMP. R, regulatory subunit; C, catalytic subunit. **B-C)** Pulldown of PKA-C-eGFP (green) captured PKA-R-mCherry (red) in the absence of cAMP (B) but not in the presence of 1 mM cpt-cAMP (C). All analyses were performed using a confocal microscope. Upper images: microbeads; middle images: magnification of boxed imaging areas; low images: corresponding surface plot maps of middle images; scale bars, 30 and 5 μm in upper and middle images, respectively. **D-E)** Statistical results of GFP (D) and mCherry (E) intensity in Panels B-C and in a similar experiments performed using HA antibody for immunoprecipitation (IP). \*\**P* = 0.0046, \**P* ≤ 0.031; n = 3 independent biological replicates, each representing the average of 7 imaging areas in the same experiment.

In another set of experiments, we tested the sensitivity of microbead-based SiMPull for pulling down proteins from cells, specifically the minimum number of cells required for the assay. GFP pulldown was again used for this test. To avoid the effect of batch-to-batch variation in transfection efficiency and accurately determine cell numbers, 100 GFP-expressing HEK293T cells were manually picked under a fluorescence microscope after dissociating the cells through trypsinization, and the cell lysate was then serially diluted at ratios representing distinct numbers of cells. In microbead-based SiMPull, GFP could be detected, with a very high S/N ratio (6.3< S/N <44), in as few as 5 GFP-expressing cells (Fig. 3E-F), and the signal intensity of GFP molecules was proportional to the number of cells even in a range as low as 5–50 cells (Fig. 3E-F). These results indicate that microbead-based SiMPull is a sensitive and quantitative assay for studying protein expression.

### Pulldown of cAMP-dependent protein kinase A (PKA) complex

The holoenzyme or complex of PKA, one of the most widely studied protein kinases, is a heterotetramer comprising a regulatory (R) subunit dimer and two catalytic (C) subunits (Fig. 4A); binding of intracellular cAMP to the R subunits results in the release the C subunits from the PKA complex (Fig. 4A). We exploited the well-characterized interaction of the R and C subunits to evaluate the performance of the microbead-based SiMPull assay for investigating PPIs.

We examined the interaction between PKA-C-eGFP and PKA-R-mCherry in HEK293T cells; the fluorescent-tags on the proteins allowed direct visualization of PKA-C and -R subunits without immunostaining and facilitated the assessment of pulldown efficiency and specificity (Fig. 4B-E) (Jain et al., 2011). When anti-GFP-coated microbeads were used in the assay, both PKA-C-eGFP and PKA-R-mCherry were pulled down; by contrast, when the microbeads used were coated with anti-HA antibody instead of anti-GFP, few GFP and mCherry molecules were pulled down (Fig. 4D-E). These results indicated that the capture of both PKA-C-eGFP and PKA-R-mCherry relied specifically on the GFP antibody. Moreover, the observed interaction between PKA-C-eGFP and PKA-R-mCherry was disrupted by the cAMP analog cpt-cAMP (Fig. 4C-E), which confirmed that the R subunit was pulled down here due to its interaction with the C subunit rather than because of nonspecific binding to anti-GFP or the microbeads.

As expected, the majority of the GFP and mCherry signals clearly colocalized (Fig. 4B), but the colocalization was incomplete; this could be due to several reasons: (1) unequal expression of PKA-C and PKA-R in HEK293T cells; (2) proper folding and fluorescence of only 75% of GFP molecules (Waldo et al., 1999); and (3) uneven quenching of the fluorescence of GFP or mCherry in a complex.

### Pulldown of TMC1 expressed in rare sensory hair cells

Sensory hair cells of the inner ear are extremely scarce, numbering only ∼3,300 per mouse cochlea (Willott et al., 2001) and ∼15,000 per human cochlea; moreover, the efficiency with which these cells can be isolated from the surrounding supporting cells is extremely low. Thus, PPIs in hair cells are currently investigated using only heterologous overexpression systems, but these are prone to artifacts and require substantiation through studies on endogenous proteins.

SiMPull has not been tested previously for its performance in the research on rare cells; here, we determined whether microbead-based SiMPull can be used for protein pulldown in sensory hair cells. We selected TMC1 as a target protein because (1) TMC1 is a component of the mechanotransduction (MT) channel in sensory hair cells and is critical physiologically and clinically (Pan et al., 2013, Pan et al. 2018); (2) excluding highly abundant proteins such as actin and prestin, most proteins expressed in hair cells, including TMC1, cannot be detected through conventional western blotting (not shown); and (3) IP performed using microbead-based SiMPull requires the target protein to be fluorescently labeled or, alternatively, to be recognized by two antibodies against distinct, non-overlapping epitopes—one for pulling down the target protein onto the microbeads, the other for identifying (immunostaining) the protein on the microbeads. TMC1 meets this last requirement because we previously generated a mouse line carrying FLAG-tagged TMC1, in which the C-terminus and N-terminus of TMC1 can be recognized by anti-FLAG antibody and a homemade anti-TMC1 antibody, respectively (Li et al., 2019).

To immunoprecipitate TMC1 from hair cells, we dissected the organ of Corti out of the cochlea—instead of using the entire cochlea—from *Tmc1*^*FLAG/FLAG*^ mice. This approach minimized the number of non-hair cells in tissue lysates and reduced the protein background-noise level because only hair cells express TMC1. TMC1-FLAG was pulled down from the cell lysate of the organ of Corti by using anti-TMC1-coated microbeads and visualized using anti-FLAG plus fluorescently labeled 2^nd^ antibody (Fig. 5). The FLAG signal was robust in the case of *Tmc1*^*FLAG/FLAG*^ mice relative to *Tmc1*^***+/+***^ mice (S/N ratio of 5.3, Fig. 5), which indicated the specificity of the FLAG signal. Moreover, when anti-TMC1 was omitted in the assay, the FLAG signal was almost eliminated, which further strengthened the conclusion that the FLAG signal represents TMC1 (Supplementary Fig. 5). Collectively, these results demonstrated that microbead-based SiMPull can be used to analyze and quantify TMC1 expression in hair cells. This is the first report of the use of SiMPull for analyzing endogenous protein expression in rare cells.

**Figure 5.**
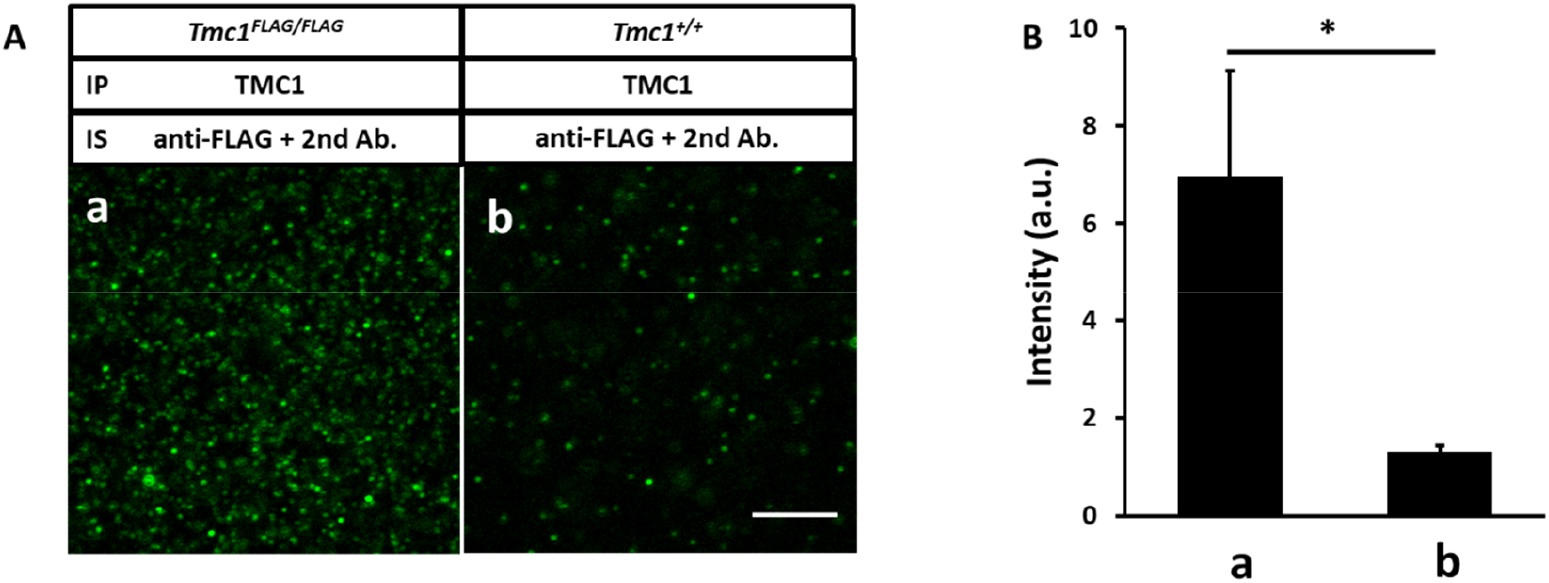
TMC1-FLAG pulldown from organ of Corti. **A)** Organs of Corti from 2–3 cochleae were dissected from *Tmc1*^*FLAG/FLAG*^ and *Tmc1*^*+/+*^ mice and lysed and homogenized, and then TMC1-FLAG in the lysates was pulled down using a homemade anti-TMC1 serum and detected using anti-FLAG plus a fluorescently labeled 2^nd^ antibody (a). *Tmc1*^***+/+***^ mice: negative control (b). IS, immunostaining; scale bar, 5 μm. **B)** Summary results of Panel A. \**P* = 0.012; n = 4 independent biological replicates, each representing the average signal intensity of 7 imaging areas in the same experiment. The S/N ratio (a/b) is 5.3.

One of the unique strengths of SiMPull is its single-molecule resolution, a feature that allows quantification of the copy number or centration of a target protein (Jain et al., 2011). Each hair cell in the assay presented in Fig. 5 was calculated to contain ∼130 fluorescent spots, each representing a functional unit of TMC1 (i.e., a TMC1 dimer or oligomer; Supplementary Information). The genuine number of TMC1 functional units per hair cells is likely to be even higher if we consider the binding efficiency of TMC1 to the beads (<100%; Supplementary Information). Nevertheless, our results indicate that the SiMPull method can provide reliable and useful information on the lower limit of the copy number (or concentration) of a target protein.

### Pulldown of LHFPL5-PCDH15 complex from sensory hair cells

Lastly, we evaluated the performance of our microbead-based SiMPull technique in the co-IP of proteins from hair cells by examining an established PPI in hair cells—the formation of a complex containing lipoma HMGIC fusion partner-like 5 (LHFPL5) and protocadherin 15 (PCDH15). LHFPL5 and PCDH15 are components of the MT complex in hair cells, and their physical interaction was recently demonstrated using cryo-EM analysis (Ge et al., 2018) in addition to the original demonstration through co-IP in heterologous expression systems (Xiong et al., 2012). We again used the dissected mouse organ of Corti to minimize the amount of non-hair cells in the samples. When LHFPL5 was immunoprecipitated from cell lysates, PCDH15 was also captured (Fig. 6); the anti-PCDH15 antibody we used was characterized previously (Schietroma et al., 2017). Conversely, little signal was detected when anti-LHFPL5 was omitted in the IP or anti-PCDH15 was omitted in the immunostaining, and the S/N ratio was ∼4 (Fig. 6). The S/N ratio is reasonably high considering that tissue samples usually have a higher background than culture cells in conventional western blotting because of complex cell types/protein compositions and potential contamination of endogenous antibodies; conceivably, the SiMPull has the same problem. Nevertheless, these results clearly demonstrated that our microbead-based SiMPull technique can be successfully used to detect PPIs in rare cells such sensory hair cells.

**Figure 6.**
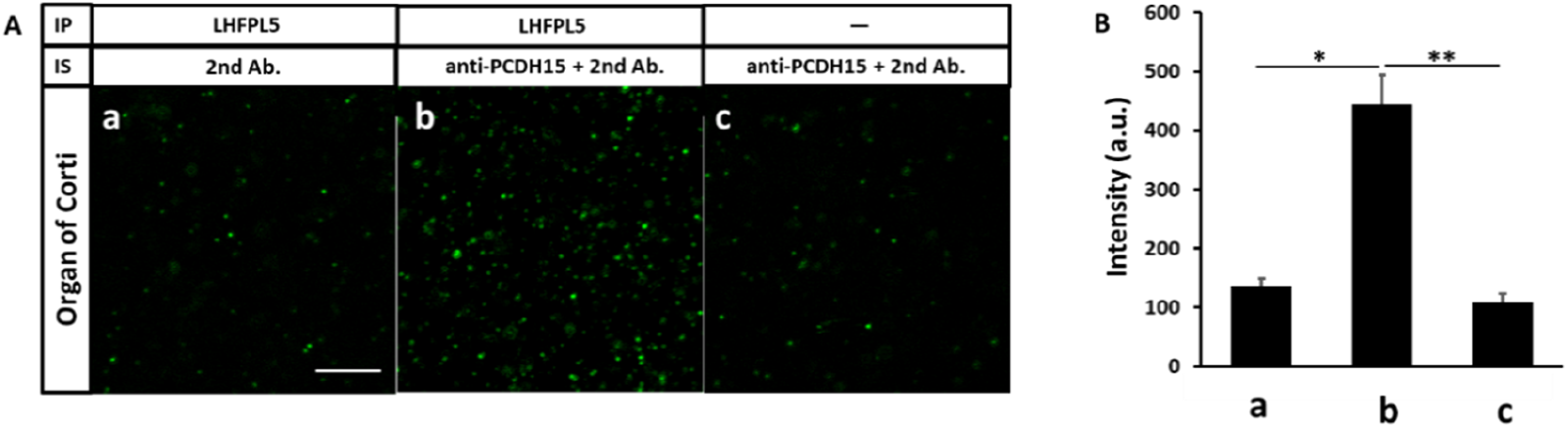
Co-IP of LHFPL5 and PCDH15 from organ of Corti. **A)** PCDH15 was captured when immunoprecipitation (IP) was performed on tissue lysates by using anti-LHFPL5. PCDH15 was detected using anti-PCDH15 plus a fluorescently labeled 2^nd^ antibody. Few PCDH15 molecules were detected when anti-PCDH15 was not included in the immunostaining (IS) (a) or anti-LHFPL5 was not used in the IP (c). Scale bar, 5 μm. **B)** Statistical results from Panel A and 2 similar experiments. \**P* = 0.011, \*\**P* = 0.0097; n = 3 independent biological replicates, each representing the average signal intensity of 7 imaging areas in the same experiment.

## DISCUSSION

The SiMPull method developed by Jain et al. has substantially expanded our capacity to analyze PPIs because the method enables evaluation of the stoichiometry of a protein complex, the kinetics of PPIs, weak and transient PPIs, and protein concentration (Jain et al., 2011). Moreover, the ultrahigh sensitivity of SiMPull allows PPIs to be analyzed in as few as 10 cells (Jain et al., 2011). However, the wide application of the original SiMPull method is greatly impeded by its relatively high technical barrier and time-consumption (Supplementary Table 1).

Because both surface-functionalization of quartz slides and the use of a micro flow-chamber are omitted in our microbead-based SiMPull, this method is considerably faster, easier to use, and more reproducible than the original SiMPull (Fig. 1 and Supplementary Table 1). Notably, the performance of our SiMPull method, in terms of sensitivity and S/N ratio, is equivalent to that of the original method (Supplementary Table 1). In principle, our SiMPull can be developed into a multiplex assay in which multiple PPIs or protein-DNA interactions are concurrently detected and analyzed by using mixed color-coded microbeads coated with antibodies; such an assay will enable side-by-side comparison of multiple PPIs and will save not only time but also analytes, which is particularly crucial in the case of rare cells or rare biological fluids. In summary, we expect the low technical barrier and high speed and efficiency of our method to considerably promote the wide application of versatile and powerful SiMPull in general research laboratories.

The simplicity of microbead-based SiMPull was exploited here in employing the technique to investigate rare sensory hair cells; this represents the first application of SiMPull in the study of rare cells. Hair cells, which number in the tens of thousands, are extremely scarce relative to the tens of millions of other sensory cells such as olfactory neurons and photoreceptors; moreover, isolating hair cells from their surrounding supporting cells is tedious and highly inefficient—an entire day’s labor results in the harvest of only a handful of hair cells (Li et al., 2019). An alternative approach that represents a compromise is to use the dissected organ of Corti or entire cochlea for biochemical analyses of hair cells; however, this method can be effectively used with western blotting (which requires ∼10^4^–10^5^ cells) only for highly abundant proteins such as actin and prestin and not for less abundant proteins, and the method cannot be used with conventional co-IP, which requires ∼10^5^–10^6^ cells, even when several cochleae are pooled (only ∼3,300 hair cells per mouse cochlea). Currently, PPIs in hair cells are examined using only heterologous overexpression systems, but the variables that affect PPIs in heterologous systems, including expression level, posttranslational modification (phosphorylation, glycosylation, etc.), and cellular context (presence or absence of certain proteins), could differ markedly from those in native hair cells. Therefore, the results obtained using heterologous systems must be substantiated through studies on endogenous proteins.

We examined TMC1 expression and analyzed the established LHFPL5-PCDH15 interaction to demonstrate that microbead-based SiMPull can be successfully used for both quantifying protein expression and investigating PPIs in hair cells (Figs. 5 and 6). Our study provides a foundation for further application of SiMPull to the study of rare cells such hair cells. Moreover, in this study, the single-molecule resolution of the SiMPull method was exploited in our quantification of the apparent number of TMC1 functional units per hair cell as 130, regardless of the oligomerization state of TMC1 (Supplementary Information); the genuine number was estimated to be at least 4 times higher because not all TMC1 molecules were pulled down and identified (Supplementary Information). Furthermore, we estimated that 30%–60% of all TMC1 molecules are present in the MT complex in stereocilia (Supplementary Information). The oligomerization state or stoichiometry of TMC1 functional units could be further investigated and validated through photobleaching analysis in the microbead-based SiMPull assay, similar to what is presented in Fig. 2, where the physical association of TMC1 subunits is more unambiguous or more physiologically relevant than that observed in previous studies (Beurg et al., 2018; Pan et al., 2018).

## ACKNOWLEDGMENTS

The work was supported by Hong Kong RGC GRF16111616, GRF16102417, and GRF16100218, NSFC-RGC joint research scheme N_HKUST614/18, Shenzhen Basic Research Scheme (JCYJ20170818114328332), SMSEGL20SC01-K (all to P.H.), and in part by the Innovation and Technology Commission (ITCPD/17-9).

## AUTHOR CONTRIBUTIONS

Conceptualization: Q.Z., Y.S., and P.H.; methodology: Q.Z., Y.S., F.T., L.Y., and H.P.; formal analysis: Q.Z. and P.H.; investigation: Q.Z., X.L., and X.Y.; resources: H.P.; writing-original and draft: Q.Z. and P.H.; writing-review and editing: P.H.; visualization: Q.Z.; supervision: P.H.; funding acquisition: P.H..

## DECLARATION OF INTERESTS

The authors declare no competing interests.

## Notes

### Competing Interest Statement

The authors have declared no competing interest.

