## Supplemental data for "Fast and easy single-molecule pulldown assay based on agarose microbeads"

**Supplementary Table 1. Features and procedures of original and microbead-based SiMPull**

| | **Original SiMPull** (Jain et al., 2011)(Jain et al., 2012) | **Microbead-based SiMPull** (this study) |
|---|---|---|
| **1. Applications** | Single-molecule imaging for interaction kinetics and stoichiometry; quantification of protein concentration; PPI of nonabundant proteins; rare cells | Same as original SiMPull |
| **2. Pulldown site** | Surface of glass coverslips | Surface of agarose microbeads |
| **3. Surface passivation & functionalization** | Required: PEG passivation (KOH/aminosilane used) and avidin coating | Not necessary: commercially available, pre-functionalized surface of microbeads |
| **4. Steps/time for 3** | 10 steps, 6–8 h | None |
| **5. Micro flow-chamber** | Required: narrow channel between coverslip and glass slide; 0.5 h needed | Not needed: all reactions performed in 1.5-mL Eppendorf tubes |
| **6. Difficulty of sample preparation** | Difficult: micropipette used to inject solution slowly into micro flow-chamber | Easy: solution added into Eppendorf tubes |
| **7. Total # of steps and time** | 21 steps<br>9–11 h | 6 steps<br>2.5–3 h |
| **8. Detection limit** | ~10 cells for detecting overexpressed proteins | ~5 cells for detecting overexpressed proteins;<br>≤ 10 pM biotin-Atto 488 |
| **9. S/N ratio** | 10–20 | 10–20 |

### Supplementary Figures

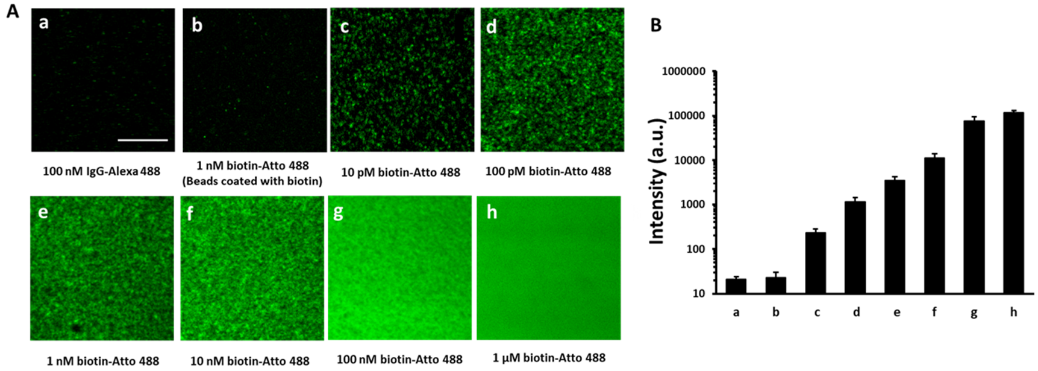


**Supplementary Fig. 1. Detection sensitivity of NeutrAvidin-coated agarose microbeads for biotin-Atto 488. A)** Biotin-Atto 488 at various concentrations pulled down by NeutrAvidin-coated agarose microbeads. In Image b, 1 nM biotin-Atto 488 was pulled down by microbeads pre-exposed to 1 μM biotin for 10 min; little binding here suggests little nonspecific binding of biotin-Atto 488 to microbeads. IgG-Alexa 488 (100 nM) is another negative control (a). Shown are 25×25 μm imaging areas selected from microbeads (see additional details in Fig. 2B and Fig. 3A-D). Scale bar, 10 μm. **B)** Average signal intensity of an imaging area similar to that in Panel A in 6 randomly selected beads in the same experiment. The microbeads detected biotin-Atto 488 at a concentration as low as 10 pM with a high signal-to-noise ratio (>10 relative to column b; the Y-axis is in the logarithmic scale).

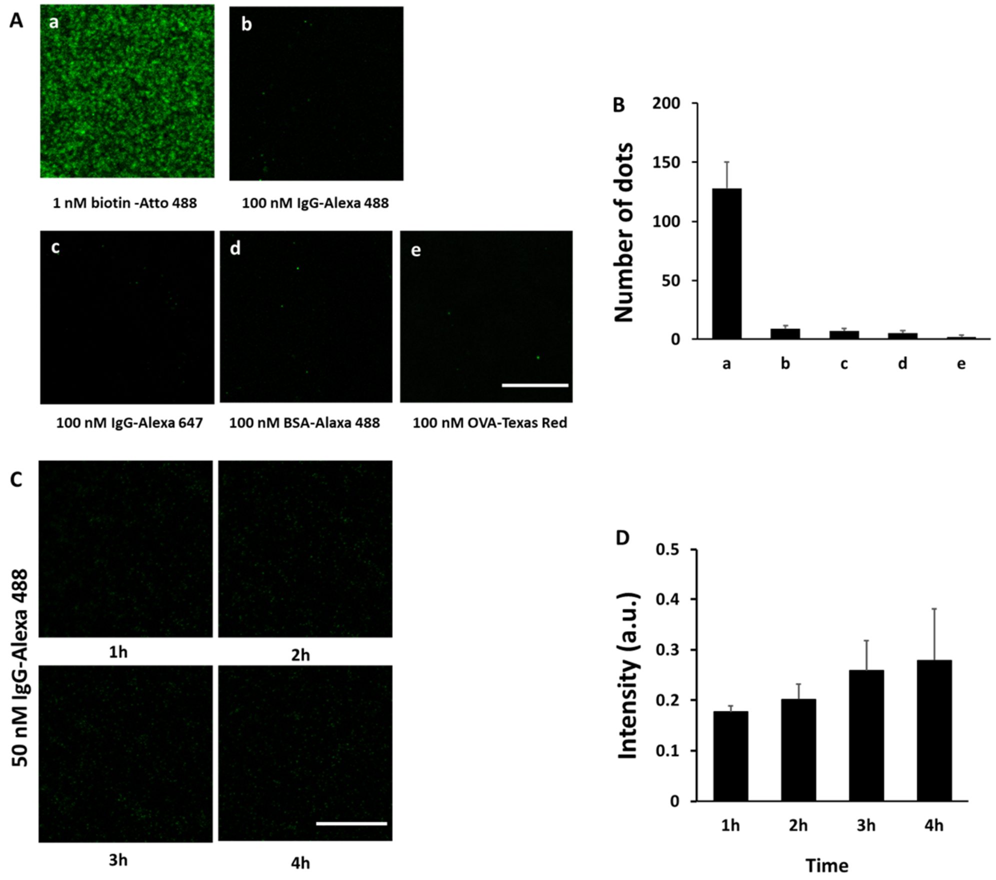


**Supplementary Fig. 2. Minimal nonspecific binding of NeutrAvidin-coated agarose microbeads. A)** Microbeads bound to biotin-Atto 488 (1 nM) as expected (a) but showed little binding, at 100-fold higher concentration (100 nM), to IgG-Alexa 488, IgG-Alexa 647, BSA-Alexa 488, or OVA-Texas Red. Shown in Panels A and C are 25×25 μm imaging areas selected from microbeads (see additional details in Fig. 2B and Fig. 3A-D); scale bars, 10 μm. **B)** Average number of fluorescent dots of an imaging area similar to that in Panel A in 6 randomly selected beads in the same experiment. **C)** Time course of nonspecific binding of 100 nM IgG-Alexa 488 to microbeads. **D)** Average signal intensity of an imaging area similar to that in Panel C in 6 randomly selected beads in the same experiment.

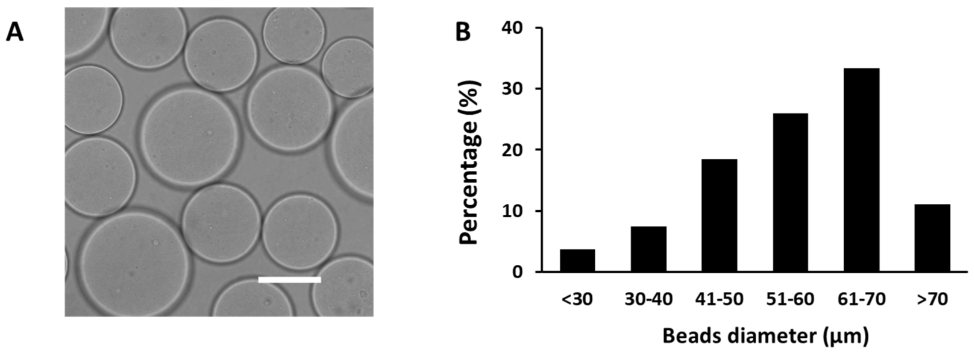


**Supplementary Fig. 3. Size distribution of NeutrAvidin-coated agarose microbeads**. A) Bright-field image of microbeads under a microscope; scale bar, 50 µm. **B)** Size distribution of a total 50 microbeads.

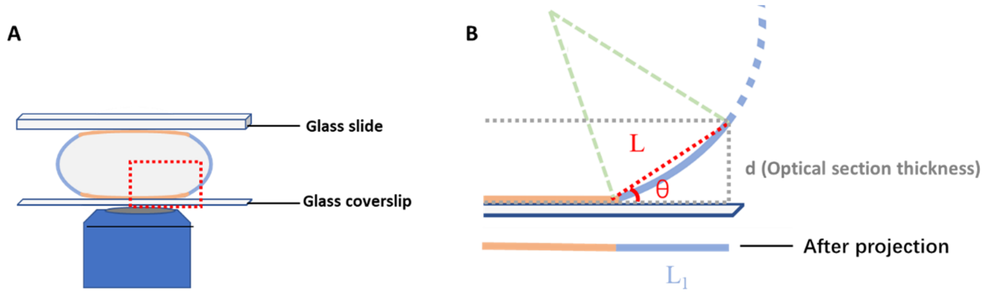


**Supplementary Fig. 4. Schematic showing formation of bright aureole on agarose microbeads**. **A)** Cross-section of a soft agarose microbead that is pressed and flattened between a coverslip and a glass slide. Blue object, microscope lens. **B)** Magnification of boxed area in panel A. The red dotted line "L" is drawn to approximate the blue curved line marking the edge of the microbead, the signal on which is projected onto Line $L_1$. Assuming that the fluorescent signal density on the bead surface is ρ, the signal density of the orange straight line that represents the central portion of the microbead remains ρ after projection; by contrast, the signal density of the curved line (approximated by L) should be $\rho(L/L_1) = \rho/\cos\theta$ after projection. Because $\cos\theta < 1$, the signal intensity of L1 is greater than ρ, and therefore the edge is brighter than the center (and an aureole is formed). This phenomenon is more readily observed under a low-magnification lens because such a lens (relative to a high-magnification lens) increases the optical section thickness d (WILSON, 2011) and concurrently increases the signal density of the aureole $\rho/\cos\theta$ (by increasing θ) and $L_1$, the width of the aureole.

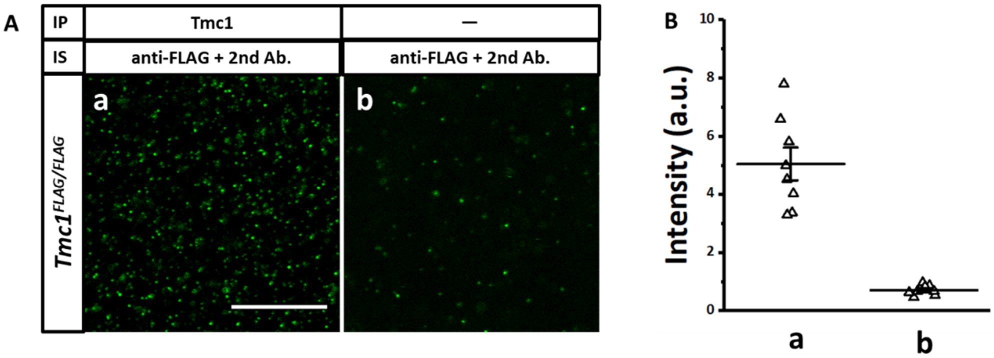


**Supplementary Fig. 5. TMC1-FLAG pulldown from organ of Corti by using anti-TMC1. A)** TMC1-FLAG was pulled down from tissue lysates by using anti-TMC1 and detected with anti-FLAG plus a fluorescently labeled 2$^{nd}$ antibody (a). Few TMC1-FLAG molecules were detected when anti-TMC1 antibody was not included in the immunoprecipitation (IP) (b, negative control). Each panel shown here is a selected (boxed) imaging area from a microbead. IS, immunostaining; scale bar, 10 μm. **B)** Statistical results of the assay in Panel A. Each triangle represents one imaging area from the experiment depicted in Panel A.

### SUPPLEMENTARY INFORMATION

#### A. Apparent number of TMC1 functional units per hair cell

The apparent number of TMC1 functional units per hair cell, N, was calculated as follows:

$$N = \frac{n_b A_b n_d}{A_i n_c}.$$

Here, $n_b$ is the number of microbeads (100 for each experiment), $A_b$ is the average surface area of a microbead (15,385 μm$^2$), $n_d$ is the number of fluorescent dots per imaging area (350 for TMC1), $A_i$ is the imaging area (25×25 μm$^2$), and $n_c$ is the number of hair cells per experiment (2 mouse cochleae, 6,600 hair cells). We assumed that each fluorescent spot represented no more than one functional unit given the low concentration of TMC1, and we obtained an N of 130 in our experiment.

The genuine number of TMC1 functional units per hair cell could be at least 4 times higher—TMC1 pulldown and identification involved 4 layers of interactions between the 1$^{st}$ and 2$^{nd}$ antibodies and TMC1, even when we exclude the extremely tight NeutrAvidin/biotinylated-2$^{nd}$-antibody interaction (see Fig. 1); this translates into only 25% of the total TMC1 molecules being captured and identified even if the efficiency at each layer of interaction is as high as 70%.

The number of TMC1 subunits in each functional unit has not been established. Dimer formation has been suggested by initial cryo-EM data and homology modeling (Ballesteros et al., 2018; Pan et al., 2018), and the existence of tetramers is another possibility (see below).

**B. Fraction of total TMC1 molecules in MT complex**

The proportion of total TMC1 molecules in the MT complex, $P_{MT}$, is calculated as follows:

$$P_{MT} = \frac{N_{MT}}{N_{total}}$$

$N_{MT}$ and $N_{total}$: number of TMC1 functional units in the MT complex of the hair bundle and in the entire cell, respectively.

The results of recent photobleaching experiments conducted by Beurg et al. suggested that a single MT complex contained 8–20 TMC1 subunits in a tonotopic gradient (Beurg et al., 2018b). The findings suggest that TMC1 could function as a dimer or tetramer if not a monomer, and this implies that each MT complex harbors an average of 7 (for dimer) or 3.5 (for tetramer) functional units. Because a single hair cell contains ~70 stereocilia in total and two-thirds of these (in the two shorter rows out of the three rows of stereocilia in total) contain a single MT complex, each hair cell contains ~45 MT complexes and therefore 315 (for tetramer) or 158 (for tetramer) TMC1 functional units, which is the value of $N_{MT}$.

$N_{total}$ is the genuine number of TMC1 functional units per hair cell, which could be 4 times the N value (see above); this yields an $N_{total}$ of 520 and thus a $P_{MT}$ of 30%–60%.
